# Fibrotic substrate stiffness enhances endometriotic epithelial cell motility

**DOI:** 10.64898/2025.12.31.697209

**Authors:** Shohini Banerjee, Nane Manukyan, Kimberly M. Stroka

## Abstract

Fibrosis is a common pathological feature of inflammatory conditions across various organ systems, leading to a marked increase in matrix stiffness. Although substrate stiffness is known to increase cell migration in cancer and stromal cell populations, it is not well understood how it affects benign epithelial cell motility, particularly within the pelvic cavity. We used an endometriotic epithelial cell line and polyacrylamide hydrogels with tunable stiffness––with glass as an extreme stiffness reference––to model mechanically driven single-cell and multicellular migration. We found that stiff substrates promoted cell speed, actin stress fiber formation, focal adhesion presentation, and spheroid expansion compared to the soft substrate. Increasing cellular contractility on the soft substrate and decreasing contractility on the stiff substrate led to an increase and decrease in cell speed, respectively. These findings provide mechanistic insight on how fibrosis as a biomechanical state regulates epithelial cell migration, with additional relevance to the pathogenesis of benign yet invasive conditions.

## INTRODUCTION

Fibrosis, tissue scarring and stiffening resulting from inflammation, occurs when fibroblasts deposit excessive amounts of extracellular matrix (ECM) components. Fibrosis is a key pathological feature of most inflammatory conditions across multiple organs.^1^ While fibrosis of the lungs, liver, heart, and kidneys are common and thus widely studied, reproductive and pelvic organs can also be affected. Conditions such as retroperitoneal fibrosis (Ormond’s Disease), endometriosis, adenomyosis, pelvic inflammatory disease, connective tissue disorders, and post-surgical trauma are all associated with inflammation leading to fibrosis of tissues in the pelvic cavity. As an excess of ECM accumulates–– primarily collagen––tissues experience a significant increase in stiffness: healthy tissues such as lung, liver, and bowel have a Young’s modulus of 1–3 kPa which increases to 16.5–20 kPa during fibrosis.^2–6^ Cells have the ability to sense and respond to mechanical stimuli such as substrate stiffness. This process of mechanotransduction results in downstream changes to cell behavior, including proliferation, migration, invasion, and epithelial-mesenchymal transition (EMT).^7,8^ These mechanically-driven cell behaviors can become dysregulated and contribute to disease severity and progression.^7,9^ For example, increased ECM stiffness promotes malignant endometrial cancer cell phenotypes via an elevated expression of ROCK1, a mechanosensitive kinase.^10^ Stiff substrates also promote actin stress fiber formation and proliferation of deep infiltrating endometriotic stromal cells.^11^ Broadly in the pelvic region, substrate stiffness has been shown to enhance the migratory and invasive phenotypes of ovarian cancer cells^12^, cervical cancer cells,^13,14^ bladder cancer cells,^15^ and colorectal cancer cells.^16,17^

The effects of substrate stiffness are well established in cancer and stromal cells, but they have not been widely studied in other benign yet invasive conditions that also exhibit fibrosis. Specifically, non-malignant epithelial mechanobiology in fibrotic microenvironments––particularly its consequences for cell motility––is underexplored. Moreover, previous studies on stiffness-mediated cell migration primarily rely on 2D cell culture models, leaving open the question of how fibrotic stiffness influences multicellular collective expansion. Thus, the mechanobiological principles governing how fibrotic substrate stiffness regulates epithelial cell motility, both at the single-cell and collective levels, remain poorly defined.

In this study, we chose to focus on an endometriotic epithelial cell type (12Z), which has relevance to benign but invasive conditions such as endometriosis. We cultured these cells on polyacrylamide (PA) gels with tunable stiffness (2 kPa as healthy; 30 kPa as fibrotic), as well as glass as an ultra-stiff extreme (∼70 GPa), to model mechanically driven cell motility within the pelvic cavity, including both 2D and 3D cell culture. Our results demonstrate a positive correlation between substrate stiffness and cell migration, mediated by cytoskeletal arrangement and Rho/ROCK signaling activity. Broadly, this work contributes to the understanding of how fibrosis as a biomechanical state regulates epithelial cell migration during the pathogenesis of benign fibrotic conditions.

## METHODS

### Cell culture

The 12Z human endometriotic epithelial cell line was purchased from Applied Biological Materials (Cat. # T0764). Cells were cultured in DMEM/F-12 (Gibco) supplemented with 10% fetal bovine serum (FBS) (Gibco) and 1% penicillin-streptomycin (Gibco). The cell line was mycoplasma free and authenticated by the company prior to shipping. Cell line authentication and mycoplasma testing were performed again at least 6 months afterwards. Cells from passages 55-80 were used for experiments.

### Polyacrylamide hydrogel preparation

Polyacrylamide (PA) hydrogels were formed on glass coverslips as previously described.^18,19^ Briefly, 22 x 22 mm glass coverslips were passed through a flame to remove contaminants before being smeared with 0.1M NaOH. Coverslips were then silanized with 3-aminopropyltriethoxysilane (Sigma Aldrich) and activated with 0.5% glutaraldehyde (Sigma Aldrich) in phosphate buffered saline (PBS) for 30 min to enable PA hydrogel attachment. 2 kPa and 30 kPa PA hydrogel precursor solutions contained certain amounts of 40% acrylamide (Bio-Rad), 2% bis-acrylamide (Bio-Rad), 1M HEPES (Gibco), and ultrapure water mixed together. Final concentrations were 4% acrylamide and 0.14% bis-acrylamide for the soft gel; 12% acrylamide and 0.28% bis-acrylamide for the stiff gel. Elastic moduli of PA gels were estimated via submersion compression testing using a Dynamical Mechanical Analyzer 850. Measurements were 1.223 ± 0.201 kPa for the soft gel (called 2 kPa in this manuscript) and 29.81 ± 2.338 kPa for the stiff gel (called 30 kPa in this manuscript). PA hydrogel precursor solutions were syringe-filtered (0.2 µm filter) and degassed for 20 min. 10% ammonium persulfate (Bio-Rad) in water and N,N,N’,N’-Tetramethylethylenediamine (TEMED; Bio-Rad) were added to the solutions to polymerize ∼80-µm-thick PA hydrogels on coverslips. PA hydrogels were allowed to polymerize for 30 min before activating with sulfo-SANPAH (Thermo Scientific) to cross-link the extracellular matrix protein to the PA hydrogel surface, as previously described.^19^ PA hydrogels were coated with 50 µg/mL collagen I from rat tail (Corning) over night at 4°C. For experiments on glass, 22 x 22 mm glass coverslips were coated in 50 µg/mL collagen I overnight at 4°C. Validation of equal coating across gels and coverslips was performed using a biotinylated collagen coating, applying avidin conjugated to Texas Red, and imaging under fluorescence microscopy. All PA hydrogels and coverslips were stored in PBS at 4°C for up to 10 days prior to cell seeding. PA hydrogels and coverslips were equilibrated in full media at 37°C at least 45 min prior to cell seeding.

### Quantification of cellular morphology

Morphological parameters including cell area, perimeter, solidity, and roundness were extracted using ImageJ to manually trace cells from phase contrast microscope images. Solidity was defined as 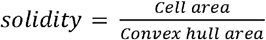, where a cell that is perfectly circular would have a value of 1.0 and a cell that has protrusions and/or indentations would have a value less than 1.0. The minor axis to major axis ratio represents the inverse of aspect ratio, where a highly elongated cell would have a value closer to 0.

### Single cell random migration

The random migration of single cells was analyzed from time-lapse phase contrast images. Briefly, approximately 80,000 cells were seeded per substrate in a 6-well plate and allowed to adhere overnight. Cells were imaged under phase contrast at 10X for 12 hr, with a time step of 15 min. Individual cell trajectories were tracked using the Manual Tracking plugin in ImageJ by clicking on the estimated geometric center of the cell at each time point. Subsequent analysis was performed in MATLAB. Average speed was defined as the cell’s total path length divided by the duration of the time-lapse. Cells were classified as fast (speed>1 µm/min), medium (0.5 µm/min<speed<1 µm/min), or slow (speed<0.5 µm/min). Directional persistence was defined as the cell’s end-to-end displacement divided by its total path length, where a cell that progresses in a perfectly straight line from point A to point B would have a directional persistence of 1; a cell that originates at a point A, moves in a straight line to point B, and directly returns to point A would have a directional persistence of 0.

### Spheroid spreading assay

Approximately 2,000-2,500 12Z cells were seeded per well in an ultra-low attachment, round-bottom 96-well plate (Corning), which was then centrifuged at 150 x*g* for 2 min. Cells self-assembled into spheroids over the next ∼48 hr of incubation. Resulting spheroids were approximately 200-250 µm in diameter. 8-10 spheroids were seeded per substrate within a 6-well plate and allowed to settle for 1 hr prior to imaging. Phase contrast imaging was then performed at 10X for 20 hr, with a time step of 20 min. The projected area of the spheroids at multiple time points was measured using ImageJ. As spheroids adhered and spread onto the substrates, the speed of the leading edge was measured using the Manual Tracking plugin in ImageJ. Specifically, each spheroid resulted in one leading edge speed which was the average of the instantaneous velocities of three circumferential points on a given spheroid: usually west, north, and east.

### Immunocytochemistry

Immunocytochemistry (ICC) was utilized to visualize cell cytoskeletal components. Briefly, cells were then fixed in 4% paraformaldehyde for 20 min on ice with gentle agitation, washed with PBS, and permeabilized in 0.2% TritonX-100 (Sigma-Aldrich) for 5 min. Samples were washed with PBS again and blocked in 2.5% goat serum (Abcam) for 1 hr at room temperature (RT). The primary antibody for paxillin was diluted in 2.5% goat serum, added to the samples, and incubated overnight at 4°C. The following day, samples were washed with PBS and incubated for 1-1.5 hr with Hoescht (1:10,000; ThermoFisher Scientific), Alexa Fluor 488 phalloidin (1:200; ThermoFisher Scientific), and the secondary antibody Alexa Fluor 488 goat anti-mouse IgG (1:200; ThermoFisher Scientific) diluted in 2.5% goat serum at RT. Samples were then washed with PBS and imaged at 40X under fluorescence using an FV3000 Laser Scanning Confocal Microscope. Fluorescence intensity across the cells was measured from segmented images in ImageJ.

### Pharmacological modulation of contractility

Y27632, an inhibitor of Rho-associated protein kinase (ROCK), was chosen to decrease cell contractility. Oleoyl-L-alpha-lysophosphatidic acid sodium salt (LPA), an activator of G protein-coupled receptors (GPCRs) that regulate cellular processes such as cell migration and mechanosensing, was chosen to indirectly increase cell contractility. Y27632 was purchased from Sigma Aldrich and dissolved in ultrapure water. 10 µM Y27632 and the vehicle, 0.1% water, were added to 12Z cells on 30 kPa substrates 30 min prior to imaging. LPA was purchased from Thermo Scientific and dissolved in 0.1% fatty acid-free bovine serum albumin (FAF-BSA; Sigma Aldrich). Final concentrations of 5 µM LPA and the vehicle, 0.0001% FAF-BSA, were added to the 12Z cells on 2 kPa substrates at the start of imaging.

### Microscopy

For all phase contrast images, live cells were imaged using the 10X objective on an Olympus IX-83 inverted microscope (Olympus, Center Valley, PA, USA) and the Olympus cellSens software. Cells were maintained at 37°C, 65% humidity, and 5% CO_2_ in an enclosed chamber surrounding the microscope.

### Statistical analysis

GraphPad Prism 10 (La Jolla, CA, USA) was utilized to generate graphs and perform statistical analyses. Outliers were identified using the ROUT method with a Q (maximum desired false discovery rate) = 1% only to remove definite outliers. Data were tested for normality. For comparisons between vehicle and E2 groups, unpaired Student’s *t*-tests were performed. For comparisons between three or more groups, ordinary one-way ANOVA with a Tukey’s multiple comparisons test was performed. Corresponding non-parametric tests were used for non-normally distributed datasets. For all tests, P values < 0.05 were considered statistically significant. Data are represented as mean ± standard error of the mean. All experiments were performed in at least triplicates.

## RESULTS

### Fibrotic substrate stiffness encourages cell spreading

12Z cells were seeded on 2 kPa, 30 kPa, or glass substrates and imaged with phase contrast microscopy the following day. Cells seeded on the softest gel and the glass appeared smaller and not as well-adhered compared to cells on the 30 kPa gel (Fig. 1A). Indeed, cells on the 30 kPa gel were significantly more spread out compared to both 2 kPa and glass substrates (Fig. 1B). However, we found no other differences in cell morphology parameters between the stiffness groups, including circularity (Fig. 1C), solidity (Fig. 1D), and minor axis:major axis ratio (Fig. 1E). Full frequency distributions for these morphological parameters are also provided (Supplementary Material S1; Fig. S1).

**Figure 1.**
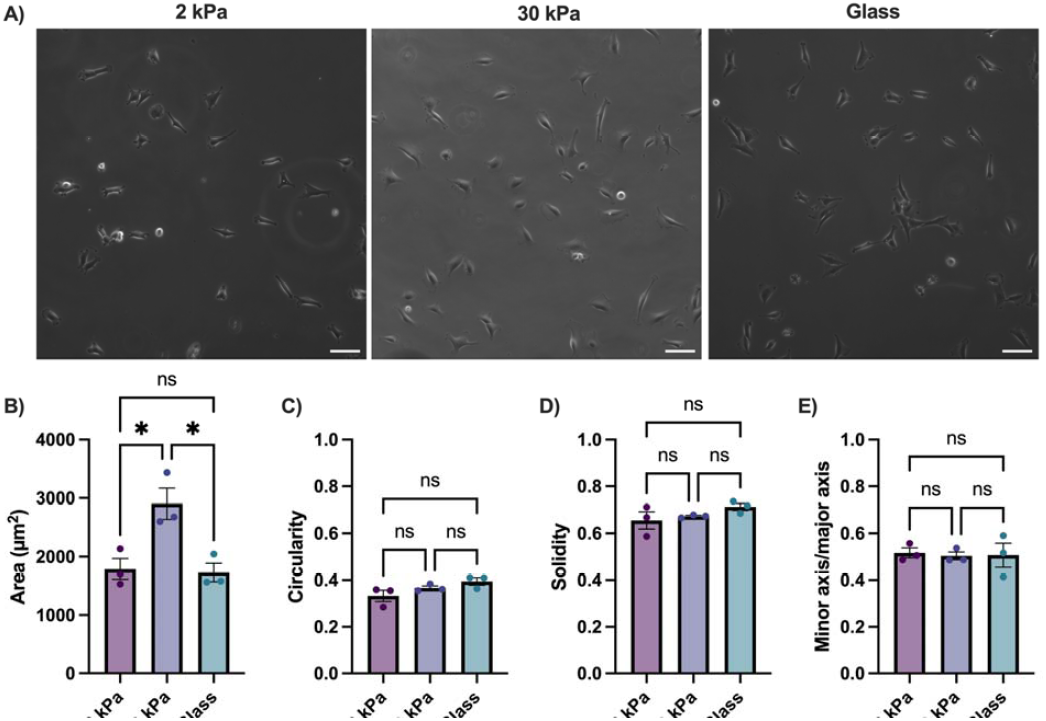
Effect of substrate stiffness on 12Z cell morphology. 12Z cells were cultured on substrates of varying stiffness overnight before assessing their morphology. **A)** Phase contrast images were taken of 12Zs on 2 kPa, 30 kPa, and glass substrates. We quantified the cell **B)** area, **C)** circularity, **D)** solidity, and **E)** ratio of the minor to major axis. Scalebars = 100 µm. Data points represent trials; error bars represent standard error of the mean. 30-40 cells were analyzed per stiffness per trial. ^*^P ≤ 0.05.

### Stiff substrates enhance single and collective cell motility

We aimed to assess the effect of substrate stiffness on endometriotic cell motility. 12Z cells were seeded on 2 kPa, 30 kPa, and glass substrates and their motility was measured the following day using timelapse microscopy. Single cell random migration was measured using Manual Tracking in ImageJ. Individual cell trajectories from the entire 12-hr timelapse show that 12Zs plated on the 30 kPa gel migrated the most extensively (Fig. 2A). The 30 kPa group contained a significantly greater proportion of fast-moving cells and a significantly reduced proportion of slow-moving cells compared to the 2 kPa and glass stiffnesses (Fig. 2B). Accordingly, cells migrated significantly faster on the 30 kPa gel compared to the other stiffnesses (Fig. 2C). Substrate stiffness did not impact cell directional persistence, defined as the end-to-end displacement divided by the total path length (Fig. 2D). For a representative subset of the cell trajectories, we plotted average cell speed over time and found that while overall speed stayed relatively steady from the beginning to the end of the 12-hr timelapse, cell speed tended to fluctuate with a period of 30-60 min across all stiffness groups (Fig. 2E). Individual cell speeds plotted over time also reflect this 30-60 min fluctuation timescale (Supplementary Material S2; Fig. S2)

**Figure 2.**
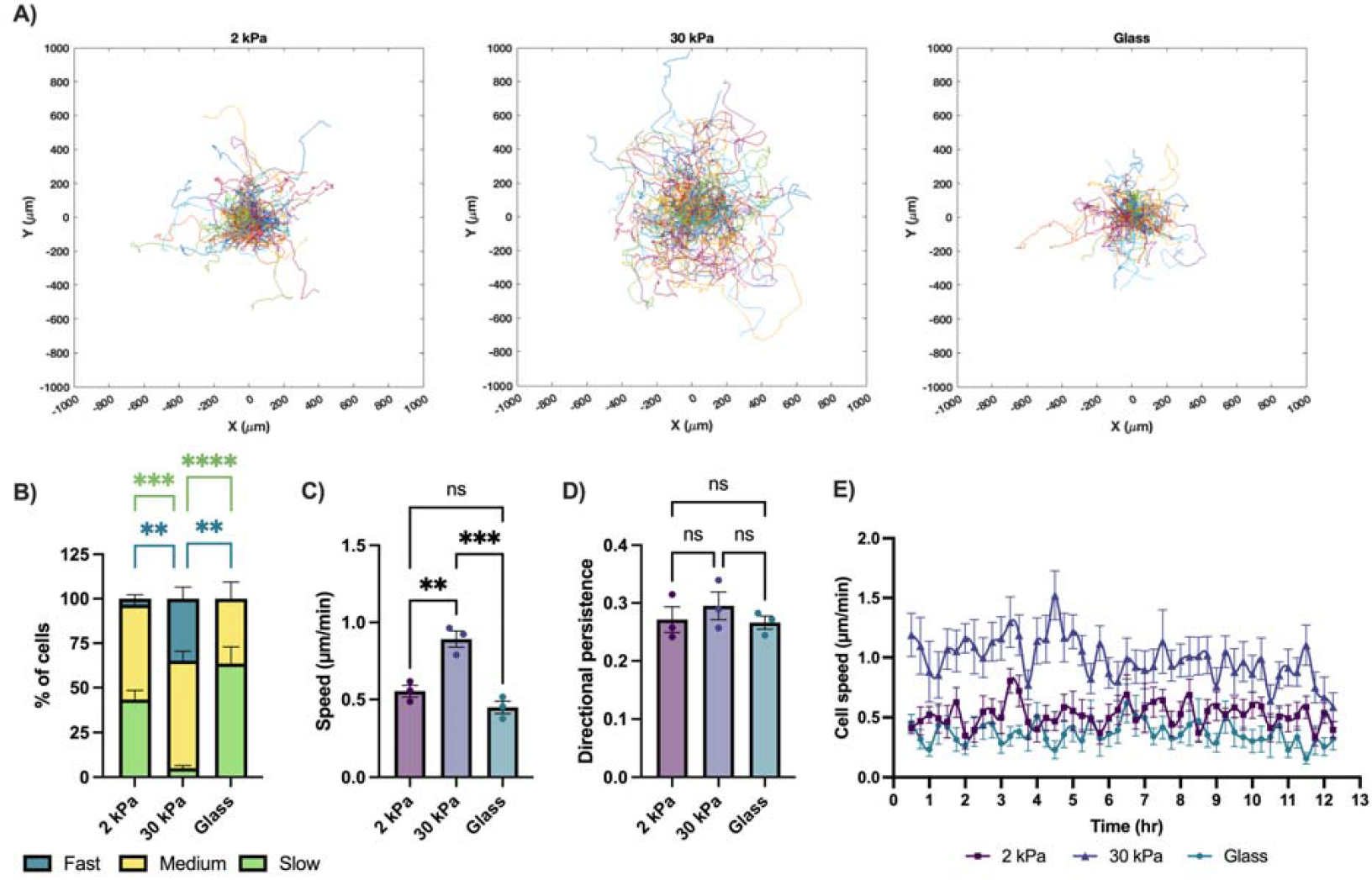
Single cell motility on different substrate stiffnesses. 12Z cells were cultured on substrates of varying stiffness overnight before measuring their random motility using timelapse microscopy for 12 hr. **A)** Trajectories of individual cells are plotted across all three trials. We quantified **B)** percentage of cells categorized as fast (speed>1 µm/min), medium (0.5 µm/min<speed<1 µm/min), or slow (speed<0.5 µm/min), **C)** average cell speed, **D)** directional persistence, and **E)** cell speed over time, where each data point represents an average from a subset of 12 cells (4 cells per trial). Unless specified, data points represent trials; error bars represent standard error of the mean. 30-40 cells were analyzed per stiffness per trial. ^**^P ≤ 0.01, ^***^P ≤ 0.001, ^****^P ≤ 0.0001.

We then aimed to study collective cell motility in a more physiologically relevant 3D model of endometriotic establishment. 12Z spheroids, approximately 250 µm in diameter, were seeded onto 2 kPa, 30 kPa, and glass substrates and allowed to settle for 1 hr prior to timelapse imaging. As cells disseminated from spheroids onto the substrate (Fig. 3A), the projected area was measured at multiple time points up to 20 hr (Fig. 3B). Spheroid outgrowth was significantly increased on the 30 kPa and glass substrates compared to the 2 kPa at 2.5 hr (Fig. 3C). A similar trend persisted at 5 hr (Fig. 3D) but the differences in outgrowth began to level out thereafter (Fig. 3B). We also measured the speed of the leading edge of the spheroids. When averaged across the entire 20-hr timelapse, there were no differences in leading edge speed (Fig. 3E); however, leading edge speed increased with substrate stiffness during early (Fig. 3F) and early-mid establishment (Fig. 3G). In this context, “early” refers to the first 2.5 hr of spheroid spreading and “early-mid” refers to the first 5 hr of spheroid spreading.

**Figure 3.**
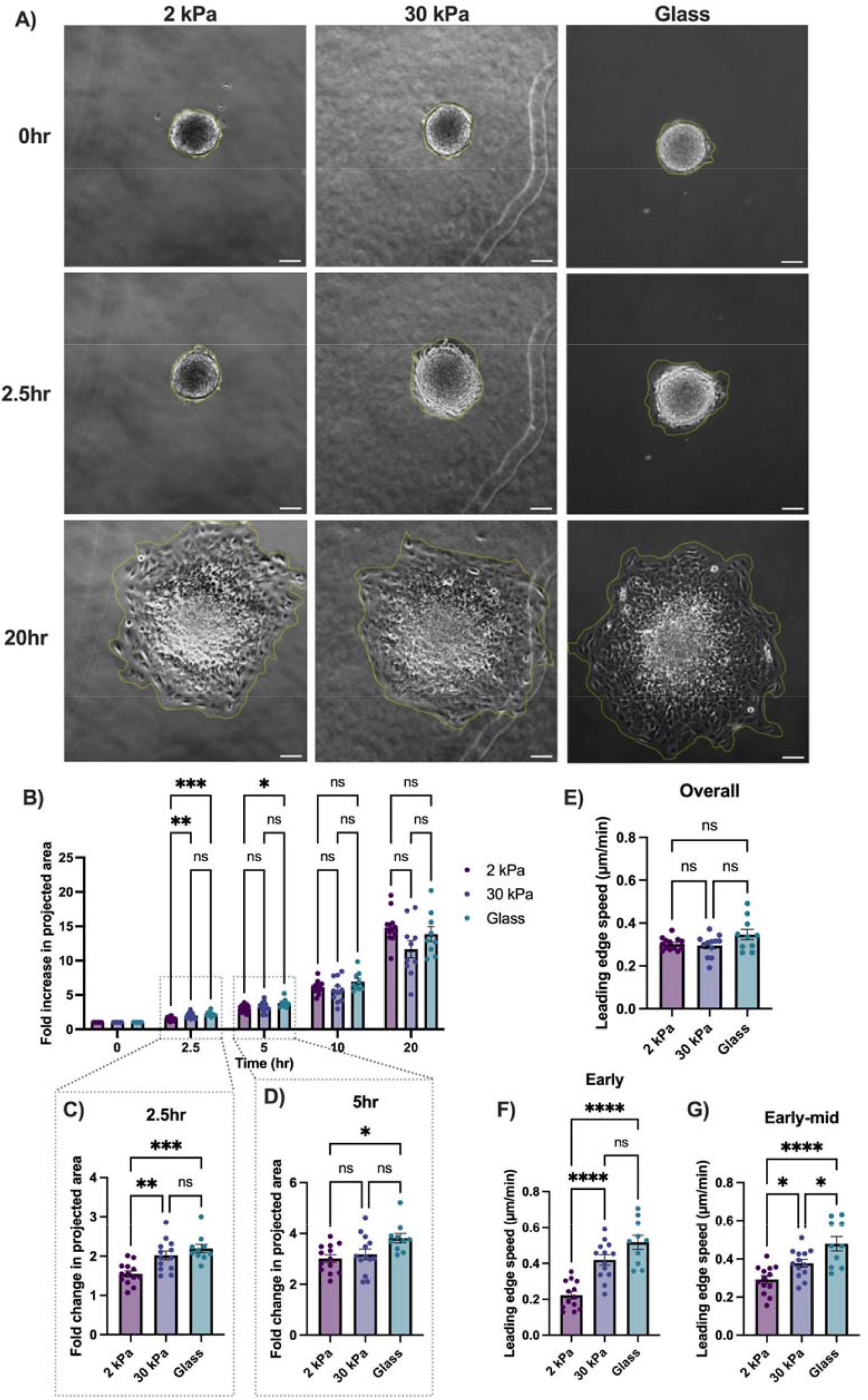
Effect of substrate stiffness on 12Z spheroid outgrowth. 12Z spheroids were seeded on substrates of varying stiffness and their outgrowth was measured using timelapse microscopy for 20 hr. **A)** Phase contrast images show spheroids on 2 kPa, 30 kPa, or glass substrates at t = 0 h, t = 2.5 hr, and t = 20 hr. We quantified **B)** the fold increase in projected spheroid area at multiple time points. We provide a zoomed in view of the fold change in projected area at **C)** t = 2.5 hr and **D)** t = 5 hr. We also measured the speed of the leading edge of the spheroid, averaging across **E)** the whole 20-hr timelapse, **F)** the first 2.5 hr, and **G)** the first 5 hr. Scalebars = 100 µm. Data points represent spheroids pooled across three trials; error bars represent standard error of the mean. 10-15 spheroids were analyzed per stiffness. ^*^P ≤ 0.05, ^**^P ≤ 0.01, ^***^P ≤ 0.001, ^****^P ≤ 0.0001.

### Substrate stiffness alters actin stress fiber presentation, membrane structures, and focal adhesions

To probe the mechanism(s) underlying differences in cell motility on different substrate stiffnesses, we analyzed cytoskeletal components in the 12Zs. Cells were plated on 2 kPa, 30 kPa, and glass substrates and fixed the following day. A fluorescent stain and ICC were used to visualize F-actin and the focal adhesion protein paxillin, respectively, in the 12Zs. Confocal images show altered actin stress fiber presentation, membrane structures, and focal adhesions across the three stiffnesses (Fig. 4A). Actin stress fibers were rarely present in 12Zs cultured on 2 kPa and glass substrates, while the majority of 12Zs on the 30 kPa substrate exhibited clear stress fibers (Fig. 4B). Of the cells that had any actin stress fibers, we observed that those on the 2 kPa gels usually had only 1-3 visible fibers, while the 30 kPa and glass exhibited more. We therefore quantified the fraction of stress fiber-presenting cells that exhibited more than 5 clear stress fibers to confirm this observation (Fig. 4C). We also observed bright spots of actin towards the periphery of cells on the 2 kPa substrates––which we call “actin puncta” ––that were much less prominent on the 30 kPa (Fig. 4D). Additionally, paxillin images show that focal adhesion density was significantly higher in cells on the 30 kPa gels than 2 kPa and glass (Fig. 4E). These focal adhesions were also larger in area (Fig. 4F).

**Figure 4.**
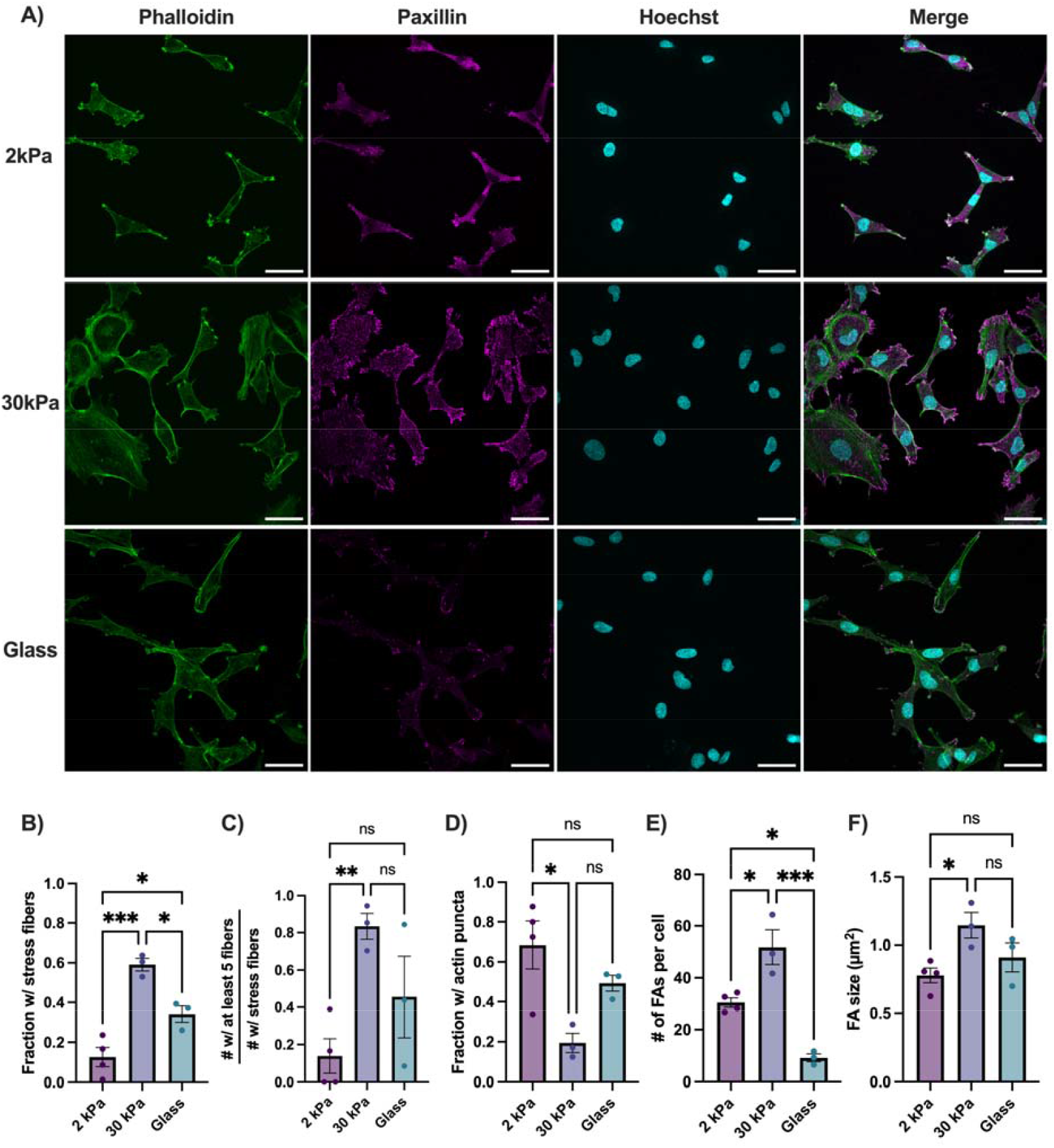
Impact of substrate stiffness on F-actin organization, membrane structures, and focal adhesions. 12Z cells were cultured on substrates overnight prior to fixation and fluorescence microscopy. Confocal images show F-actin (Phalloidin), focal adhesion proteins (paxillin), and nuclei (Hoechst) for cells on 2 kPa, 30 kPa, and glass substrates. We quantified **B)** the fraction of cells exhibiting actin stress fibers, and **C)** the fraction of stress fibered cells exhibiting more than 5 stress fibers, **D)** the fraction of cells exhibiting punctate actin membrane structures, **E)** the average number of focal adhesions per cell, and **F)** the focal adhesion size. Scalebars = 50 µm. Data points represent trials; error bars represent standard error of the mean. 4-5 images were analyzed per stiffness per trial. ^*^P ≤ 0.05, ^**^P ≤ 0.01, ^***^P ≤ 0.001. FA; focal adhesion.

### Pharmacological modulation of contractility alters cell motility on soft and stiff substrates

To further validate our finding that substrate stiffness impacts cell motility due to alterations in cell cytoskeletal components (Fig. 4) we reassessed 12Z motility with pharmacological modulation of cellular contractility. Cells were seeded on 2 kPa and 30 kPa gels and treated and imaged the following day. LPA– –an activator of GPCRs that activate pathways such as Rho/ROCK, PI3K/Akt, and MAPK/ERK––was applied to cells on the 2 kPa gels at a 5 µM dose to increase contractility. The selective ROCK inhibitor Y27632 was applied to cells on the 30 kPa gels at a concentration of 10 µM to decrease contractility. Samples were only imaged for 4 hr due to the propensity of LPA to bind plastic and thus get depleted in the well. Phase contrast images of cells treated for ∼2 hr revealed that 30 kPa-grown cells were more spread out on the substrate compared to 2 kPa, consistent with our earlier finding (Fig. 1B), and that 30 kPa cells treated with Y27632 had a distinctly shrunken appearance (Fig. 5A). LPA increased the random migration of 12Zs on the 2 kPa gels while Y27632 decreased migration on the 30 kPa gels, as seen in the single-cell trajectories (Fig. 5B). Accordingly, 2 kPa cell speed was significantly increased with LPA treatment while 30 kPa cell speed was significantly reduced with Y27632 treatment (Fig. 5C). The contractility modulators had no effect on directional persistence of cells (Fig. 5D). Similar to our result in Fig. 2E, cell speed fluctuated over time with a period of 30-60 min across all groups (Fig. 5E).

**Figure 5.**
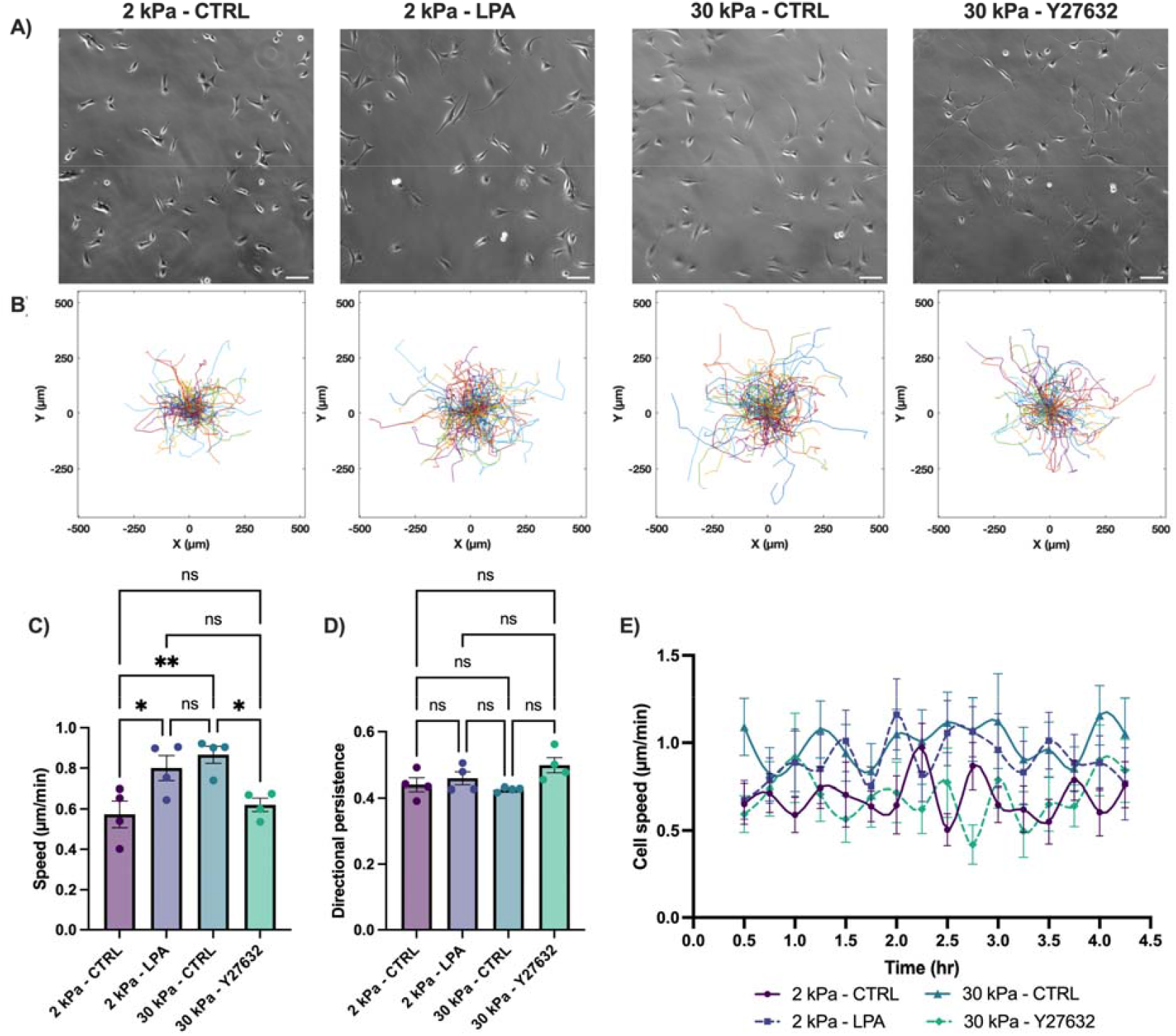
Role of cell contractility in substrate stiffness-mediated cell motility. 12Z cells were cultured on 2 kPa and 30 kPa substrates overnight before contractility modulators––LPA and Y27632, respectively––were added and cell motility was measured via timelapse microscopy for 4 hr. **A)** Phase contrast images show 12Zs cultured on 2 kPa and 30 kPa after ∼2 hr of treatment with the vehicle or drug. **B)** Trajectories of individual cells are plotted across all trials. We quantified **C)** average cell speed, **D)** directional persistence, and **E)** cell speed over time, where each data point represents an average from a subset of 12 cells (3 cells per trial). Scalebars = 100 µm. Unless specified, data points represent trials; error bars represent standard error of the mean. 30-40 cells were analyzed per stiffness per trial. ^*^P ≤ 0.05, ^**^P ≤ 0.01. CTRL; vehicle control. LPA; lysophosphatidic acid.

## DISCUSSION

In this study, we aimed to study the role of fibrotic tissue stiffening in non-malignant epithelial cell motility. Our data show that fibrotic substrate stiffness increases the migratory behavior of endometriotic epithelial cells for both single-cell and multicellular spheroids. 12Zs cultured on the 30 kPa PA gel demonstrated greater cell spreading area, increased speed, prominent actin stress fibers, enhanced focal adhesions, and greater early spheroid spreading compared to cells on the soft 2 kPa gel. These data suggest that fibrotic tissues may enhance cell migration by optimizing cytoskeletal architecture to generate contractile forces.

However, it is worth noting that for the most part, substrate stiffness did not enhance migratory cell phenotypes without limit. While cells migrated faster and had larger area on 30 kPa gel compared to those on the soft gel, cell speed and area was significantly reduced on glass compared to 30 kPa. This bell-curve shaped trend persisted with cytoskeletal arrangement as well: actin stress fibers and focal adhesions were most pronounced on the 30 kPa gel, with reduced paxillin and stress fibers on the 2 kPa and glass substrates. These findings are consistent with other key mechanobiological studies that show that migratory cell phenotypes are impaired on extremely soft and extremely stiff substrates, and that optimal migration occurs on a goldilocks stiffness or adhesion strength in between.^19–22^ Our only finding that did not reflect this phenomenon was that both 30 kPa and glass promoted early (2.5-5 hr) 12Z spheroid expansion, with glass exhibiting the greatest outgrowth and leading edge speed. In fact, spheroids on glass exhibited a significantly faster leading edge compared to both 30 kPa and 2 kPa during the first 5 hr of expansion. This early outgrowth on glass suggests that collective migration may rely on different force-generation mechanisms than single-cell migration, potentially involving population-level tension or cooperative adhesion dynamics that are less sensitive to substrate compliance.

Pharmacological modulation of contractility further supported a mechanistic role for cytoskeletal force generation in stiffness-driven cell migration. Increasing contractility and migration pathways in cells on the 2 kPa gel led to an increase in speed, approximately equal to the speed of cells cultured on the 30 kPa gel. Conversely, cells on the 30 kPa gel experienced a reduction in speed, approximately equal to the speed of cells on the 2 kPa gel, when treated with a contractility inhibitor. These results indicate that substrate stiffness tunes the level of intracellular tension required for efficient epithelial migration and are consistent with findings in other mechanobiological studies that modulated contractility.^20,23–26^

Limitations and caveats of this work must be addressed. We only utilized a single cell line, and while the 12Zs are a useful model for epithelial mechanobiology, more cell lines should be tested as different epithelial subtypes may exhibit distinct mechanosensitivities. Spheroid outgrowth in 3D could not be assessed due to the cytotoxic nature of PA gels; future work could embed 12Z spheroid in collagen gels of increasing stiffnesses to study mechanically driven invasion in 3D. Third, the use of LPA to enhance contractility, while informative, affects multiple signaling pathways and therefore does not isolate a mechanotransduction pathway. Future studies could focus on dissecting the role of specific signaling pathways in stiffness-mediated cell motility. This work creates other avenues for future research; the small actin puncta we observed frequently in cells on the 2 kPa and glass substrates could be further characterized. We speculate that they may represent points of adhesion to the substrate, since cells cannot spread fully and have minimal contact with the substrate. Lastly, the mechanobiology of epithelial monolayers––how epithelial sheet dynamics are affected by substrate stiffness––would be worth exploring.

Despite the limitations, the data identify robust and reproducible mechanobiological principles governing how epithelial cells reorganize their cytoskeleton and migrate within fibrotic mechanical microenvironments. Understanding how substrate stiffness regulates multiscale epithelial cell motility may shed light on the pathogenesis of benign but invasive fibrotic conditions and potential therapeutic targets.

## AUTHOR CONTRIBUTIONS

KMS was the principal investigator. SB contributed to experimental design, data collection, data analysis, data interpretation, and manuscript preparation. NM contributed to data collection and data analysis. All authors critically reviewed the manuscript.

