## Supplementary Figure S1 for "Fibrotic substrate stiffness enhances endometriotic epithelial cell motility"

**SUPPLEMENTARY MATERIAL S1**

To assess the effect of substrate stiffness on cell morphology, we seeded 12Z cells on 2 kPa, 30 kPa, and glass substrates and allowed them to adhere overnight before imaging. Morphology parameters were derived from tracing cells in ImageJ and distributions of cell area (Fig. S1A), circularity (Fig. S1B), minor axis/major axis ratio (Fig. S1C), and solidity (Fig. S1D) are shown.

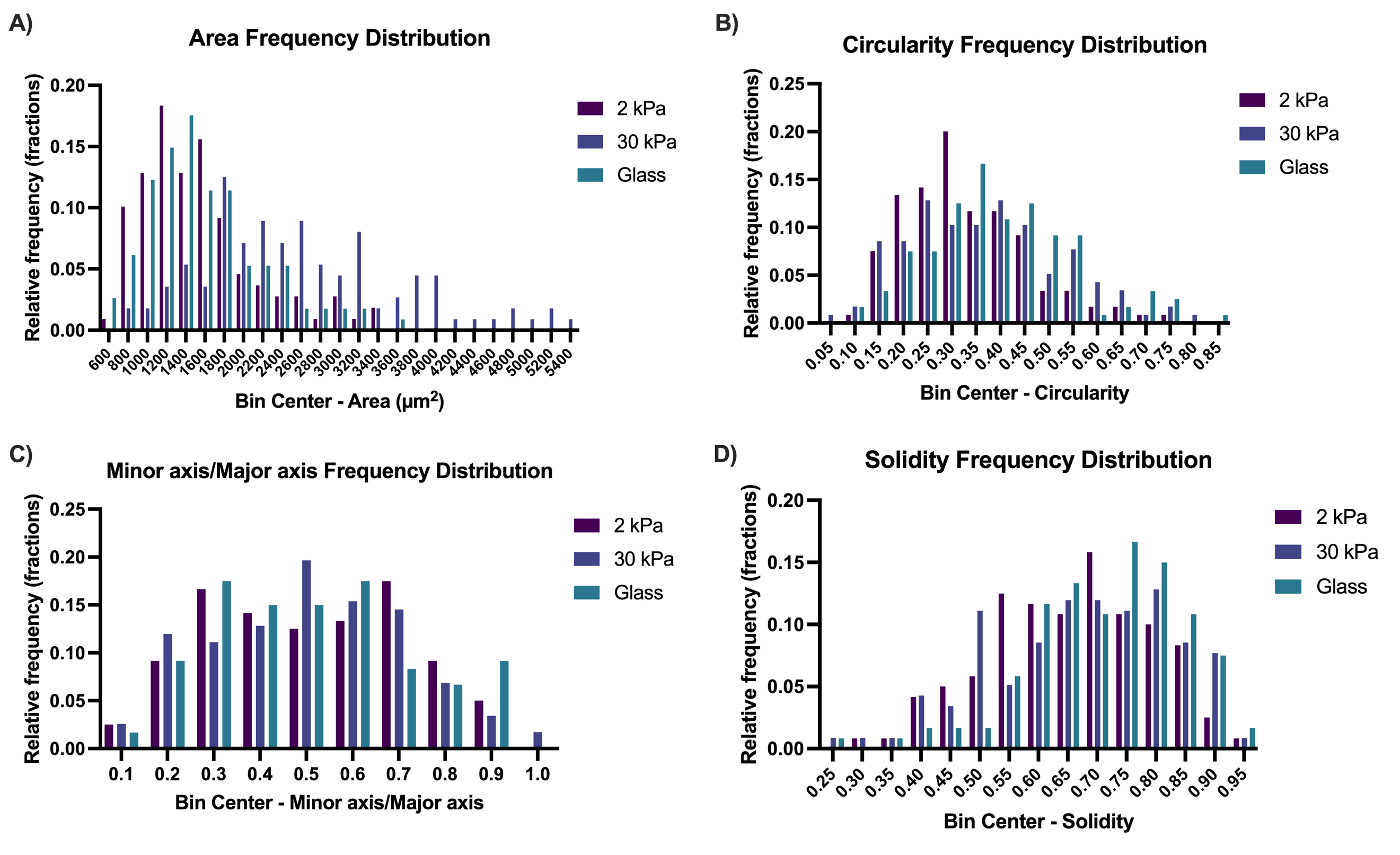
**Figure S1. Frequency distributions of 12Z morphological parameters on varying substrate stiffnesses.** 12Z cells were cultured on substrates of varying stiffness overnight before assessing their morphology from phase contrast images. Relative frequency distributions of cell morphological parameters, including **A)** area, **B)** circularity, **C)** minor to major axis ratio, and **D)** solidity were plotted and include data from all three trials. 30-40 cells were analyzed per stiffness per trial.
