## Supplementary Figure S2 for "Fibrotic substrate stiffness enhances endometriotic epithelial cell motility"

**SUPPLEMENTARY MATERIAL S2**

To assess the effect of substrate stiffness on endometriotic cell migration, 12Z cells were seeded on 2 kPa, 30 kPa, and glass substrates and their motility was measured the following day using timelapse microscopy. Single cell random migration was measured using Manual Tracking in ImageJ. For a representative subset of the cells analyzed, we plotted individual cell speed over time and found that while overall speeds stayed relatively steady from the beginning to the end of the 12-hr timelapse, they tended to fluctuate with a period of 30-60 min across all stiffness groups (Fig. S2).

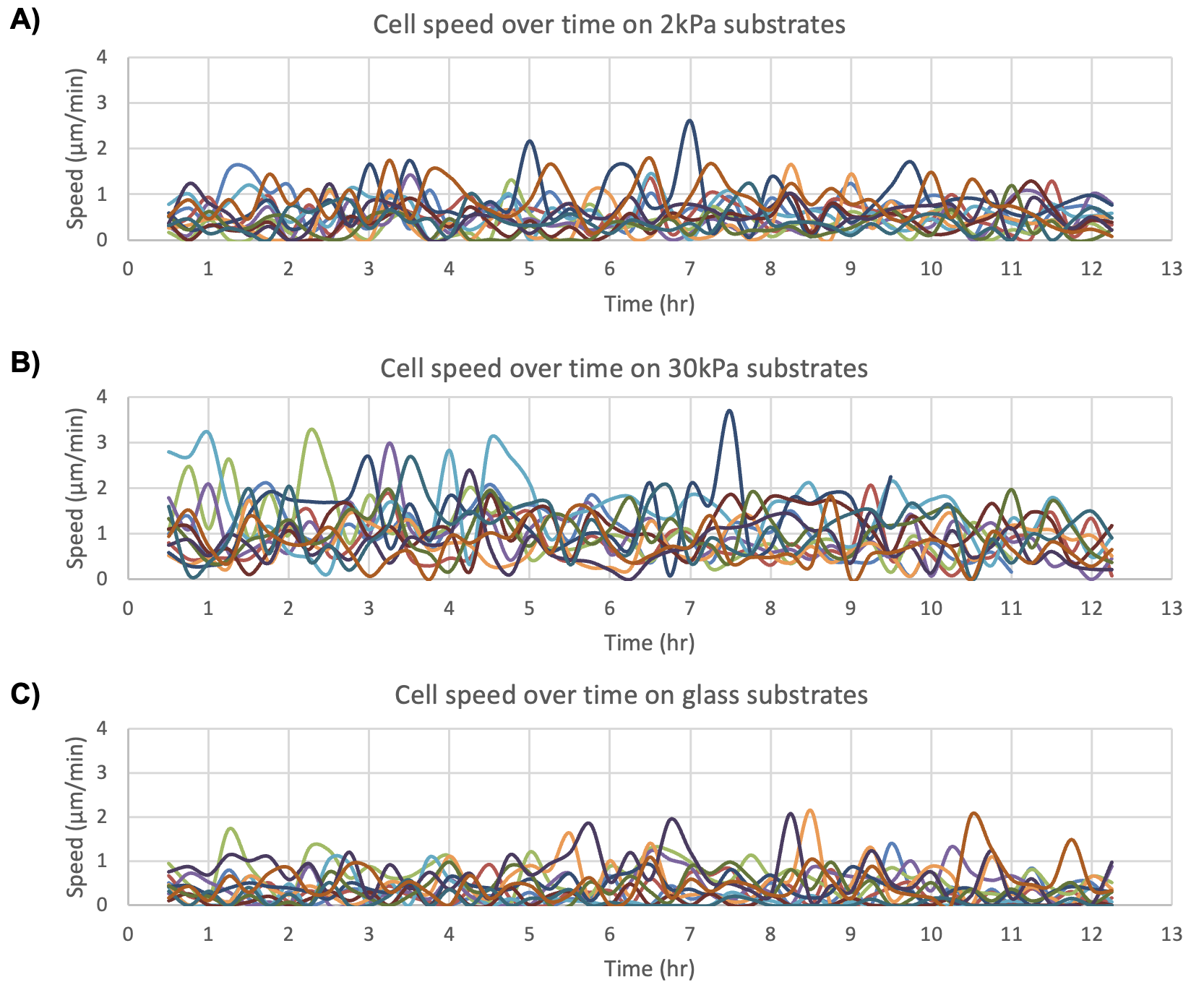

**Figure S2. Instantaneous cell speed over time on varying substrate stiffnesses.** 12Z cells were cultured on substrates of varying stiffness overnight before assessing their motility from 12-hr phase contrast timelapses. Cell speed over time is plotted for **A)** 2 kPa, **B)** 30 kPa, and **C)** glass substrates, where each curve represents a cell and each plot contains a subset of 12 cells (4 cells per trial). Instantaneous cell speed was calculated at every 15 mins.
